# SAD-1 kinase controls presynaptic phase separation by relieving SYD-2/Liprin-α autoinhibition

**DOI:** 10.1101/2023.06.12.544643

**Authors:** Nathan A. McDonald, Li Tao, Meng-Qiu Dong, Kang Shen

## Abstract

Neuronal development orchestrates the formation of an enormous number of synapses that connect the nervous system. In developing presynapses, the core active zone structure has been found to assemble through a liquid-liquid phase separation. Here, we find that the phase separation of SYD-2/Liprin-α, a key active zone scaffold, is controlled by phosphorylation. Using phosphoproteomics, we identify the SAD-1 kinase to phosphorylate SYD-2 and a number of other substrates. Presynaptic assembly is impaired in *sad-1* mutants and increased by overactivation of SAD-1. We determine SAD-1 phosphorylation of SYD-2 at three sites is critical to activate its phase separation. Mechanistically, phosphorylation relieves a binding interaction between two folded SYD-2 domains that inhibits phase separation by an intrinsically disordered region. We find synaptic cell adhesion molecules localize SAD-1 to nascent synapses upstream of active zone formation. We conclude that SAD-1 phosphorylates SYD-2 at developing synapses, enabling its phase separation and active zone assembly.

## Introduction

Developing neurons extend polarized axons and dendrites great distances to make connections with partner cells and wire the nervous system. As partners are identified, synaptic junctions are formed that will support neuronal communication. Each synapse builds specialized pre- and post-synaptic machinery to support extremely rapid asymmetric communication across the junction^1^. In postsynapses, postsynaptic densities cluster receptors in order to receive and propagate incoming neurotransmitter signals. In presynapses, synaptic vesicles containing neurotransmitters are clustered and primed for rapid release upon incoming action potential signals^2^.

The central structure of a presynapse is the “active zone”, an electron-dense membrane-apposed structure marking the site of release of synaptic vesicles^2, 3^. The active zone comprises multiple large multi-valent scaffolding proteins, including Liprin-α, RIM, RIM-BP, Piccolo/Bassoon, ELKS, and Munc-13^2^. These molecules form the active zone structure and coordinate the central functions of the presynapse, including the clustering of voltage-gated calcium channels and tethering and priming of synaptic vesicles.

While the molecular composition of the active zone is well established, how this structure develops and the signaling that initiates its formation is not clear^4^. Many binding interactions have been identified that link the various active zone scaffold molecules^2, 3^, suggesting their assembly into a densely bound matrix at nascent synapses. Recent evidence additionally suggests multiple active zone components are capable of liquid-liquid phase separation (LLPS) to form condensates. LLPS is a mechanism where multi-valent, low-affinity interactions lead to demixing of proteins or nucleic acids into dense, but still fluid, condensates^5, 6^. RIM and RIM-BP were first identified to form condensates *in vitro*^7^. These condensates were competent to cluster voltage-gated calcium channels, a key function of RIM and RIM-BP, and possess plausible interactions with synaptic vesicles *in vitro*^8^. We recently showed *C. elegans* SYD-2/Liprin-α and ELKS also formed condensates, and that *in vivo*, SYD-2 and ELKS phase separation activity was critical for active zone assembly^9^. SYD-2 and ELKS condensates acted to robustly assemble active zone components during a transient liquid state during synaptogenesis. Correspondingly, homologous mammalian Liprin-αs have been shown to phase separate, with Liprin-α3 phase separation linked to active zone structure^10^.

Beyond the presynaptic active zone, synapses contain additional condensate compartments. Synapsin and Synaptophysin form condensates on synaptic vesicles and contribute to synaptic vesicle clustering^11, 12^. Condensates have also been observed at sites of ultrafast endocytosis next to the active zone^13^. Further, in postsynapses, multiple components including PSD-95, SynGAP^14^, and Rapsyn^15^ have been identified to phase separate. These mounting observations indicate phase separation is a common compartmentalization mechanism in synapses and is critical for their formation^16^ and function^17^.

The mounting evidence for phase separation in synapses is contrasted with how little is known of the regulation of synaptic phase separation *in vivo*. It is presumably critical to form pre- and post-synaptic condensates at the appropriate developing synaptic junction site and regulate their properties to achieve a functional structure. Here, we identify phosphorylation as a mechanism controlling SYD-2/Liprin-α phase separation and presynaptic active zone formation. We identify the SAD-1 kinase to phosphorylate SYD-2 and determine mechanistically how this phosphorylation regulates phase separation. We find SAD-1 is localized to nascent synapses to activate presynaptic assembly during development through SYD-2 phase separation.

## Results

### Phosphorylation regulates SYD-2 phase separation and presynapse formation

Post-translational modifications, including phosphorylation, regulate a variety of phase-separated condensates in cells^18^. To determine if phosphorylation could be regulating presynaptic active zone condensates formed by SYD-2/Liprin-α, we first immunoprecipitated endogenous GFP-SYD-2 from *C. elegans* and probed for serine and threonine phosphorylation. We find SYD-2 is indeed phosphorylated (Figure 1A), consistent with proteome-wide phosphorylation datasets^19^. We sought to test if SYD-2’s phosphorylation regulates its phase separation and subsequent functions in presynapse assembly. We endogenously fused a promiscuous lambda phosphatase domain (λpptase) to SYD-2’s C-terminus to constitutively dephosphorylate the protein. This fusion tag effectively removed phosphorylation from SYD-2 (Figure 1B), while fusion with a catalytically inactive λpptase^H76N^ did not change SYD-2’s native phosphorylated state. With this λpptase allele, we tested the impact of dephosphorylation on the formation and liquidity of SYD-2 condensates with fluorescence recovery after photobleaching (FRAP) assays at nascent embryonic synapses (Figure 1C), where SYD-2 has been shown to be in a liquid condensate state^9^. Wildtype SYD-2 at nascent synapses recovers quickly after photobleaching, due to liquid condensate formation and rapid exchange between condensate and cytoplasmic pools^9^. These dynamics are lost when SYD-2’s intrinsically disordered region (IDR) responsible for phase separation is removed^9^. The SYD-2-λpptase fusion showed a similar phenotype, with a decrease in FRAP dynamics demonstrating inhibition of liquid condensate formation (Figure 1C-D).

**Figure 1.**
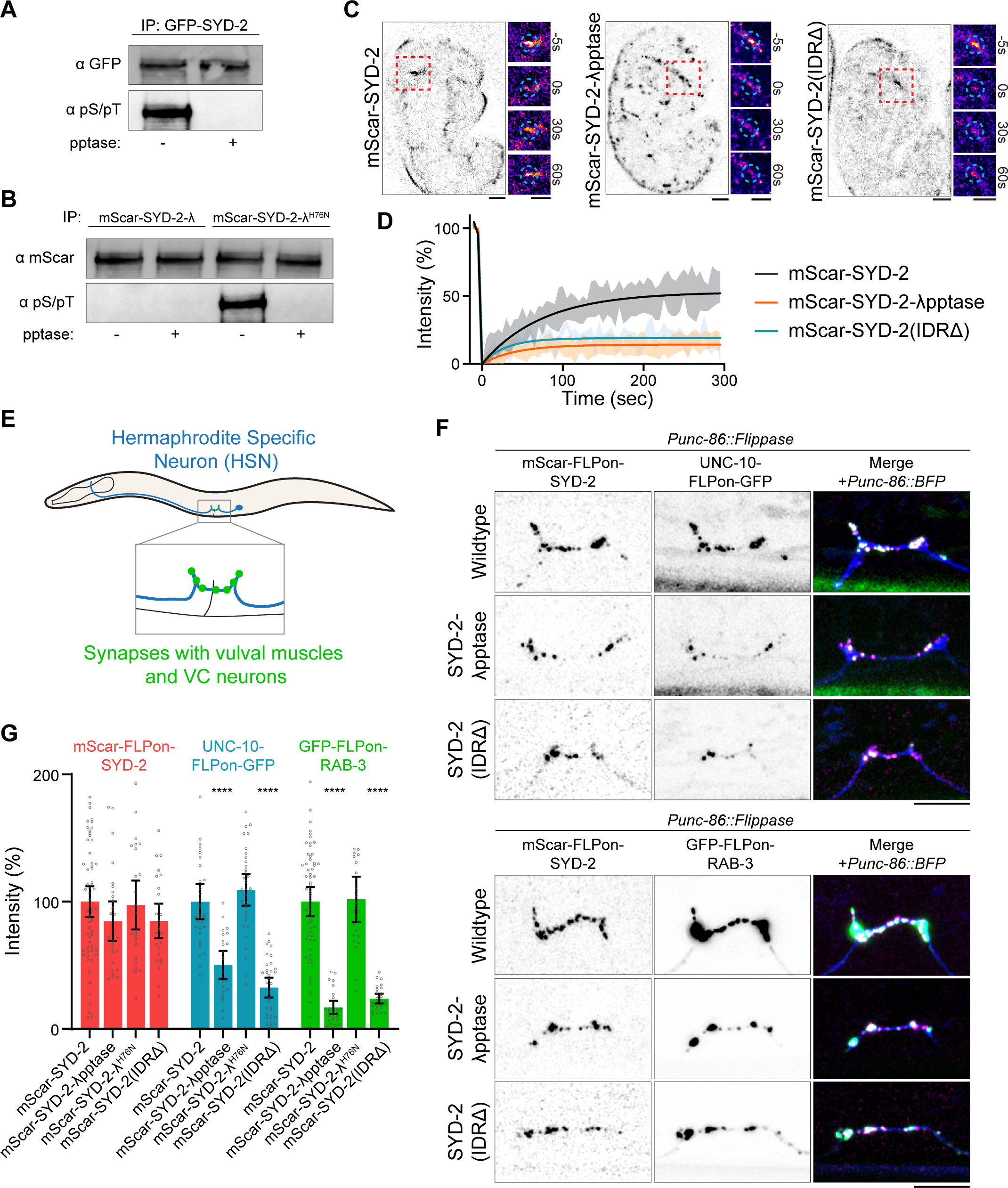
Phosphorylation regulates SYD-2/Liprin-α’s phase separation and active zone formation. A-B) Immunoprecipitation of endogenous GFP-SYD-2 (A) and endogenous mScarlet-SYD-2-λpptase (B). Western blot for GFP or mScarlet and phosphoserine/phosphothreonine. Select immunoprecipitated samples were treated with phosphatase (pptase +). C) Fluorescence recovery after photobleaching of endogenous mScarlet-SYD-2 at embryonic nerve ring synapses to measure nascent synapse condensate formation. Removal of phase separation in SYD-2(IDRΔ) or dephosphorylation in SYD-2-λpptase inhibits condensate formation. Scale bars, 1 µm. D) Quantification of FRAP in (C). E) Schematic of the *C. elegans* Hermaphrodite Specific Neuron (HSN) which makes stereotyped synapses to vulval muscles to control egg laying. F) HSN synapse formation phenotypes visualized with endogenous fluorescent tags in the indicated mutants. Scale bars, 5 µm. G) Quantification of HSN intensity in (F). ****, p<0.0001.

To next determine the impact of phosphorylation on SYD-2’s functions in presynapse formation that depend on phase separation, we imaged synapse formation in the *C. elegans* Hermaphrodite-Specific Neuron (HSN), which makes specific stereotyped synapses to vulval muscles in order to control egg laying^20, 21^ (Figure 1E). We imaged cell-specific, endogenously tagged UNC-10/RIM, a downstream core active zone component^22^, and RAB-3, a synaptic vesicle marker^23^. The SYD-2-λpptase fusion caused a reduction in synaptic UNC-10 and failed to build large RAB-3 synaptic vesicle pools (Figure 1F-G). Intriguingly, SYD-2 levels remained normal at these synapses – a phenotype consistent with a complete loss of phase separation mutant, SYD-2(IDRΔ), which localizes normally but is unable to build robust synapses (Figure 1F-G). A H76N catalytically inactive λpptase fusion had no impact on SYD-2 function or synapse formation, indicating that the fusion with λpptase does not itself perturb SYD-2 (Figure 1G). These phenotypes therefore suggest phosphorylation of SYD-2 is critical to enable its phase separation and build a presynapse.

### The SAD-1 kinase phosphorylates SYD-2 to activate phase separation and presynapse assembly

A variety of kinases and signaling pathways have been implicated in synapse formation^24^. One candidate that stood out for regulation of SYD-2 is the SAD-1 kinase, which has been linked to neuronal development in vertebrates^25–27^ and *C. elegans*^28–30^. When we imaged HSN synapse formation in a *sad-1Δ* mutant, we find a similar phenotype to SYD-2(IDRΔ) or SYD-2-λpptase alleles (Figure 2A-B). SYD-2 localizes normally, but downstream UNC-10 and synaptic vesicle accumulation are reduced. In addition, we find that overexpression of constitutively active SAD-1(T202E) increases the accumulation of SYD-2, downstream active zone components, and synaptic vesicle marker RAB-3 at synapses (Figure 2A-B and Figure S2). SAD-1(T202E) overexpression was performed with an *egl-6* promoter, chosen to achieve delayed expression in HSN to avoid major polarization and axon guidance defects associated with earlier overexpression^29^. These data argue that *sad-1* is a key regulator of presynapse accumulation during synaptogenesis. Further, these synapse formation phenotypes are consistent with a role in activating SYD-2 phase separation. Indeed, in a *sad-1Δ* mutant, SYD-2 condensate formation at nascent embryonic synapses was inhibited, as determined by FRAP (Figure 2C-D). We endogenously tagged SAD-1 with GFP and imaged its localization in HSN and find SAD-1 localizes distinctly to presynaptic sites (Figure 2E), positioning it appropriately to regulate presynapse formation and SYD-2. These phenotypes suggest SAD-1 signaling, either directly or indirectly, could be responsible for SYD-2 condensate phosphoregulation.

**Figure 2.**
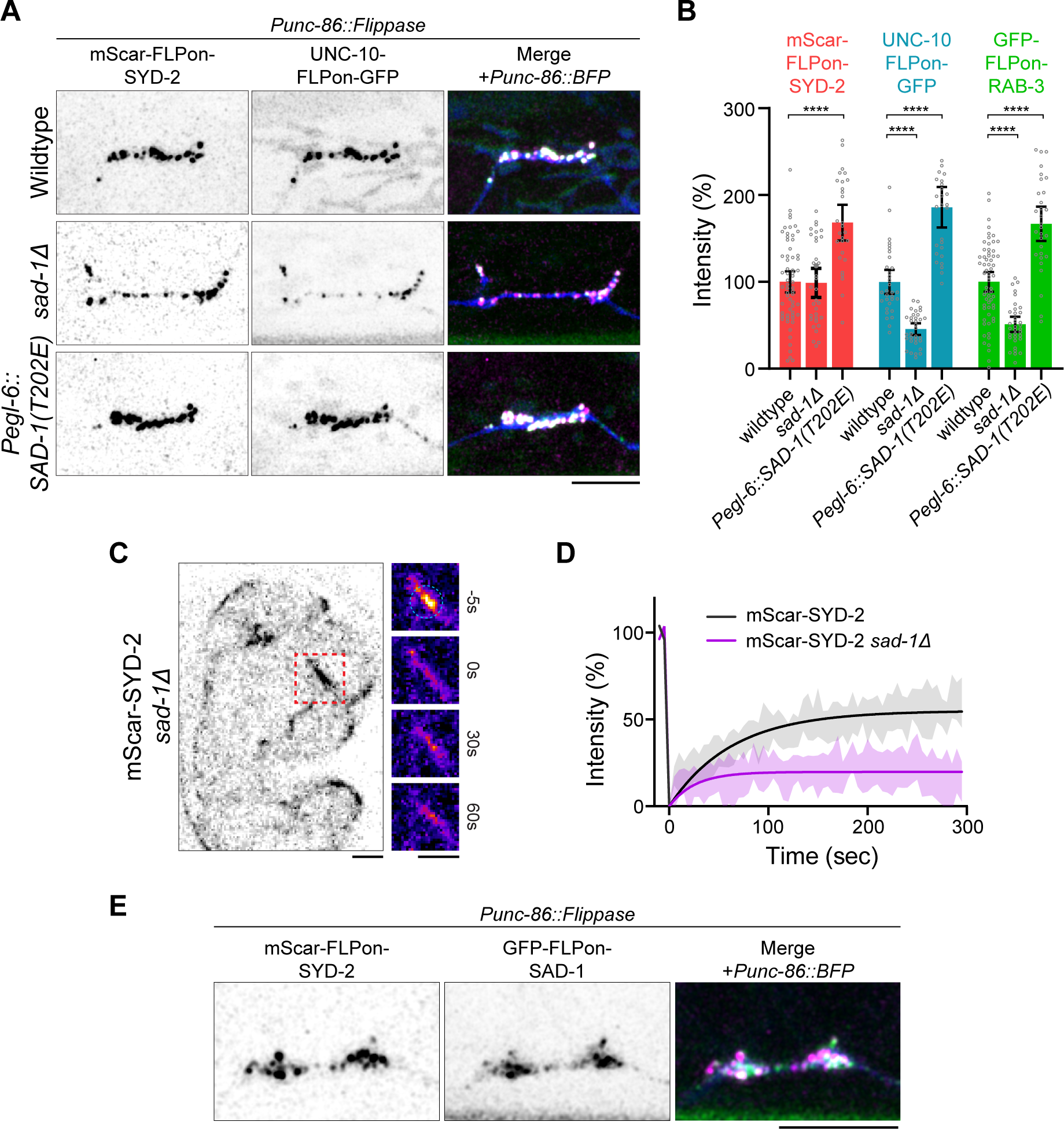
The SAD-1 kinase regulates SYD-2/Liprin-α phase separation and active zone formation. A-B) HSN synapse formation phenotypes visualized with endogenous fluorescent tags in the indicated mutants. *Pegl-6::SAD-1(T202E)* is the overexpression of constitutively active SAD-1 kinase in HSN. Scale bars, 5 µm. B) Quantification of HSN intensity in (A). ****, p<0.0001. Wildtype data is from Figure 1F. C) Fluorescence recovery after photobleaching of mScarlet-SYD-2 in a *sad-1Δ* background at embryonic nerve ring synapses to measure nascent synapse phase separation liquidity. Scale bars, 1 µm. D) Quantification of FRAP in (C). E) Endogenous localization of SAD-1 at HSN presynapses. Scale bar, 5 µm.

Despite SAD-1’s implication in synapse formation and neuronal polarity, its substrates to accomplish these functions are not known. To identify possible substrates of SAD-1, we performed a phosphoproteomics screen (Figure 3A). Wildtype animals were labeled with heavy ^15^N and mixed with *sad-1Δ* ^14^N animals, before lysis, protein digestion, and phosphopeptide enrichment. Phosphopeptides were identified and quantified by LC-MS/MS and those with high ^15^N/^14^N ratios represented potential SAD-1 substrates (Figure 3A). A variety of candidate substrates were discovered that may be phosphorylated by SAD-1 *in vivo* (Figure 3A and Table S1). Fortuitously, a SYD-2 phosphopeptide was a top hit present in all 3 biological replicates (Figure 3A), indicating SAD-1 may be responsible for SYD-2 phosphoregulation.

**Figure 3.**
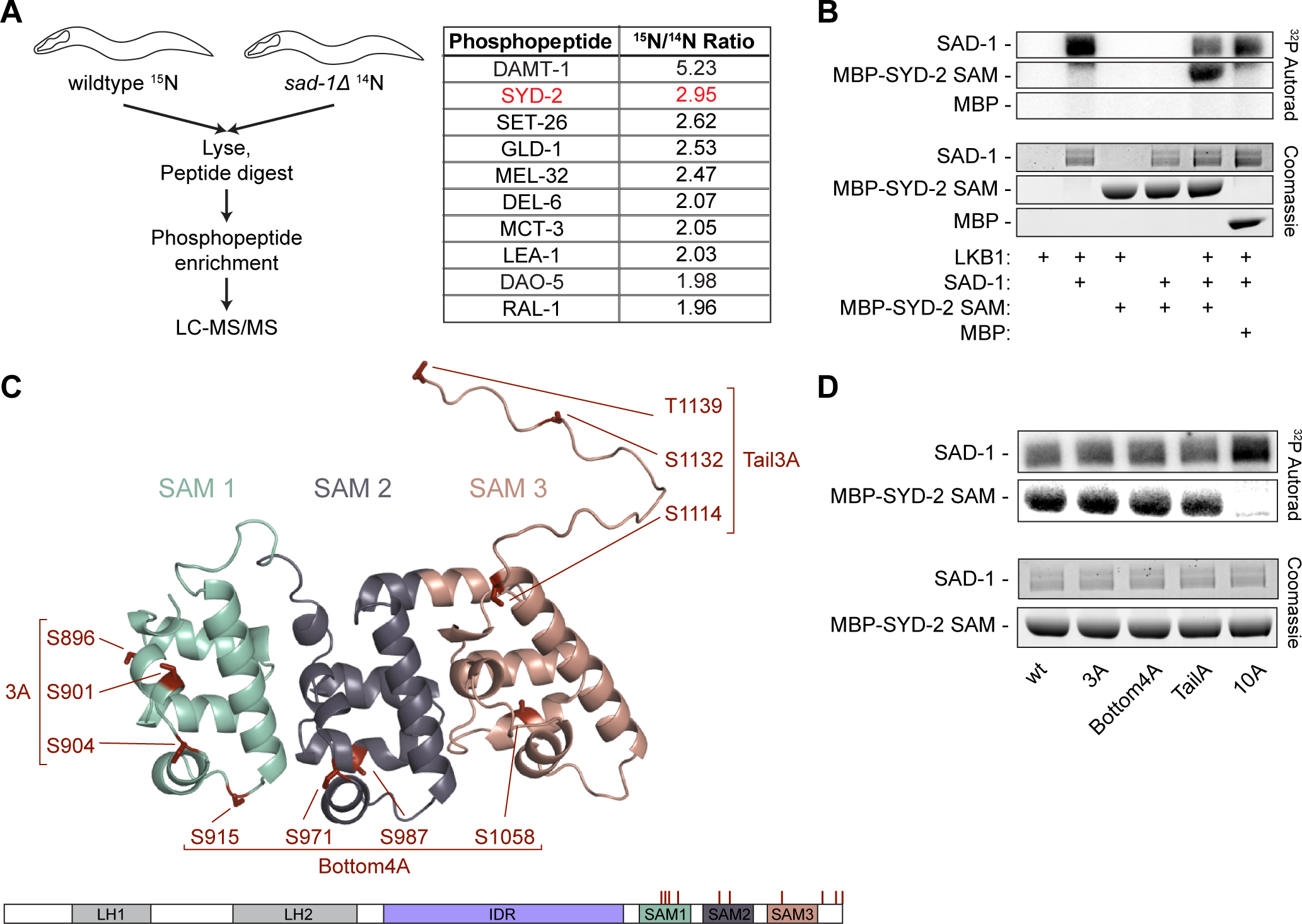
SAD-1 kinase phosphoproteomics reveal direct phosphorylation of SYD-2. A) *In vivo* SAD-1 phosphoproteomics experiment to identify possible substrates. Protein names for the top ten phosphopeptides identified are shown. See Table S1 for complete data. B) *In vitro* kinase assay between SAD-1 and SYD-2’s C-terminal SAM domains. SAD-1 is activated by the LKB-1 kinase complex. C) Alphafold 2 model of SYD-2’s SAM domains, residues 863-1139. Potential phosphosites identified from the phosphoproteomics in (A) and *in vitro* kinase assay in (B) are shown. Candidate phosphosites are organized into groups based on location for follow-up testing. See also Figure S3A and Tables S1-2. D) *In vitro* kinase assay between SAD-1 and SYD-2 phosphomutants. SAD-1 is activated by the LKB-1 kinase complex.

To test if SAD-1 was directly responsible for phosphorylating SYD-2 and determine if additional sites may be present not found in the phosphoproteomics screen, we performed *in vitro* kinase assays between SAD-1 and SYD-2 (Figure 3B and S3A-B). Purified recombinant SAD-1 kinase was pre-activated through phosphorylation by the LKB-1 complex^28^ and mixed with purified SYD-2 fragments. Activated SAD-1 was effective at phosphorylating SYD-2’s SAM and IDR domains *in vitro* (Figure 3B and S3A-B). We mapped sites from these assays with mass spectrometry, identifying 10 sites on SYD-2’s SAM domains and 26 sites in SYD-2’s IDR (Figure 3C and S3A).

We anticipated that not all of the sites identified from *in vitro* phosphorylation assays would represent true *in vivo* phosphosites, as *in vitro* kinase conditions are generally promiscuous. Indeed, only a few phosphosites were seen in our *in vivo* phosphoproteomics or in previous whole-proteome datasets (Figure S3A)^19^. *In vitro*, however, these sites were robust and only mutation of all 10 sites in the SAM domains (Figure 3D) or 26 sites in the IDR (Figure S3B) was capable of abolishing phosphorylation. We therefore separated sites into groups based on structural location to evaluate *in vivo* function (Figure 3C and S3A). We introduced alanine mutations at each group of sites into the endogenous SYD-2 gene and imaged HSN synapse formation to assay function. The 26 sites in SYD-2’s IDR, including 12 sites clustered in a key phase separation motif^9^, showed no evidence of presynaptic functionality, with normal synapse formation phenotypes in HSN (Figure S3C-D). IDR phosphomutants also showed no synaptic transmission defects in an assay for cholinergic synaptic transmission (Figure S3E). We conclude IDR phosphorylation by SAD-1 is not required for SYD-2’s function and may be an artifact of *in vitro* phosphorylation conditions. However, a 3A (S896A, S901A, S904A) mutation of sites clustered in the most N-terminal SAM domain showed a significant synapse formation phenotype (Figure 4A-B and S4), similar to SYD-2-λpptase or *sad-1Δ*, with diminished UNC-10 and RAB-3 assembly and normal SYD-2 levels. Other clusters of sites on SYD-2’s SAM domains did not impact synapse formation. A complete 10A mutant reproduced the synapse formation defects of the 3A mutant, but also impacted SYD-2 overall levels at synapses, perhaps due to impacts on protein stability. Consistently, the 3A phosphomutant also showed synaptic transmission defects, similar to the complete loss of phase separation IDRΔ mutant (Figure 4C), indicating synapses are not formed or functioning normally in the absence of this phosphorylation.

**Figure 4.**
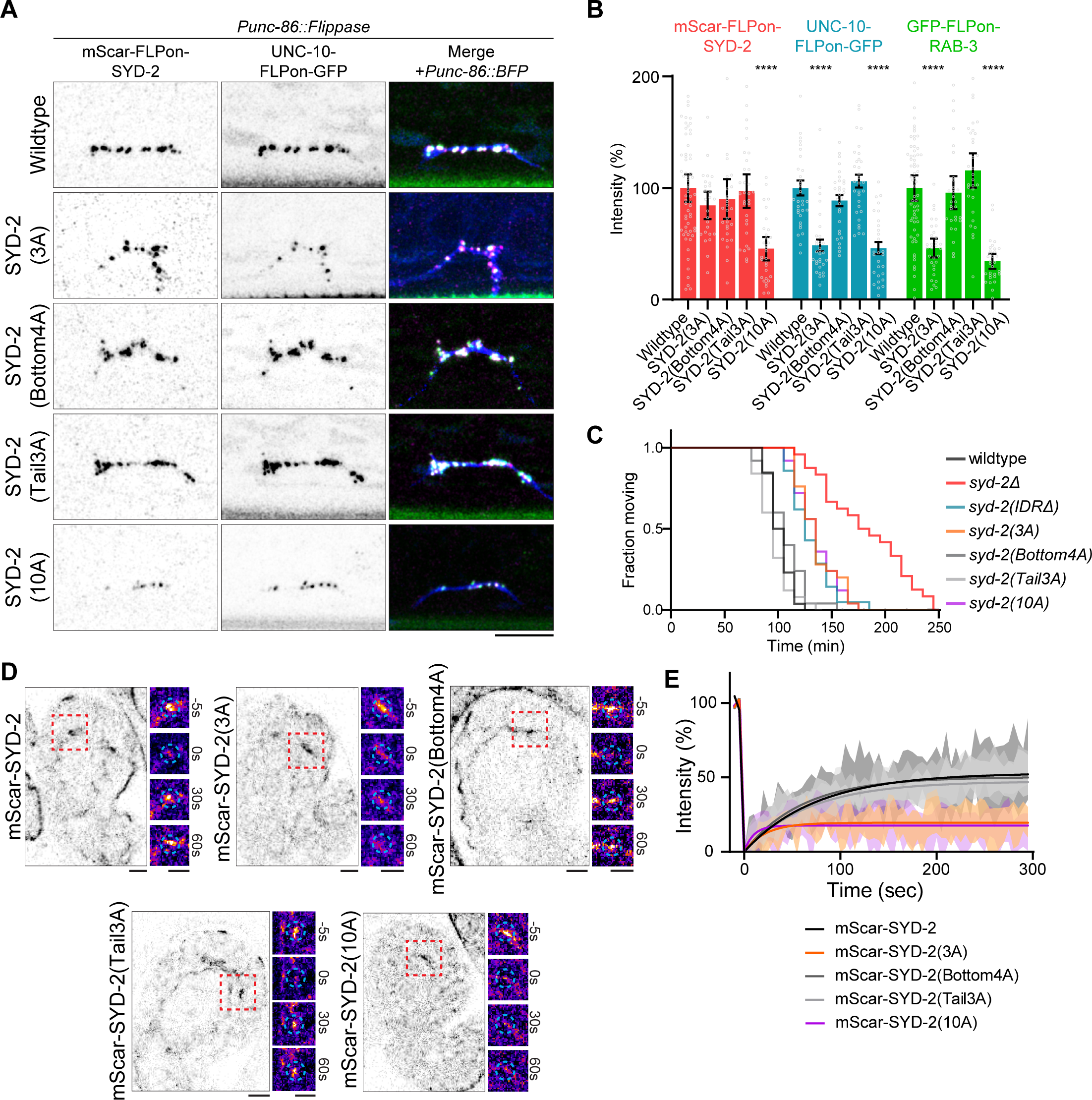
SAD-1 phosphorylation of SYD-2 controls phase separation and active zone assembly. A) HSN synapse formation phenotypes visualized with endogenous fluorescent tags in the indicated mutants. Scale bars, 5 µm. B) Quantification of HSN intensities in (A). ****, p < 0.0001. C) Aldicarb synaptic transmission assay. Additional time to paralysis on 1 mM Aldicarb indicates defective synaptic transmission. n > 20 for each genotype. D) Fluorescence recovery after photobleaching of mScarlet-SYD-2 phosphomutants at embryonic nerve ring synapses to measure nascent synapse phase separation liquidity. Scale bars, 1 µm. E) Quantification of FRAP in (D). Wildtype data from Figure 1C.

To test whether these phenotypes result from the regulation of SYD-2’s phase separation, we assayed SYD-2 phosphomutants for condensate formation at developing synapses with FRAP. The 3A and 10A mutants showed significant loss of dynamics (Figure 4D-E), indicating inhibition of condensate formation, while other SAM mutation clusters had no effect.

These phenotypes suggest three sites in SYD-2’s SAM domains are critical for regulating its phase separation and presynaptic function. To determine if these SYD-2 phosphosites account for SAD-1’s function in promoting synapse formation, we extended the overexpression of constitutively active SAD-1(T202E) into a SYD-2(3A) background. In the 3A phosphomutant, SAD-1 was not able to increase levels of SYD-2, UNC-10, and synaptic vesicles at HSN presynapses (Figure 5A-B) as seen in a wildtype background. Together, these results indicate SAD-1 phosphorylates three important sites within SYD-2’s SAM domain to enable SYD-2 phase separation and presynaptic active zone formation.

**Figure 5.**
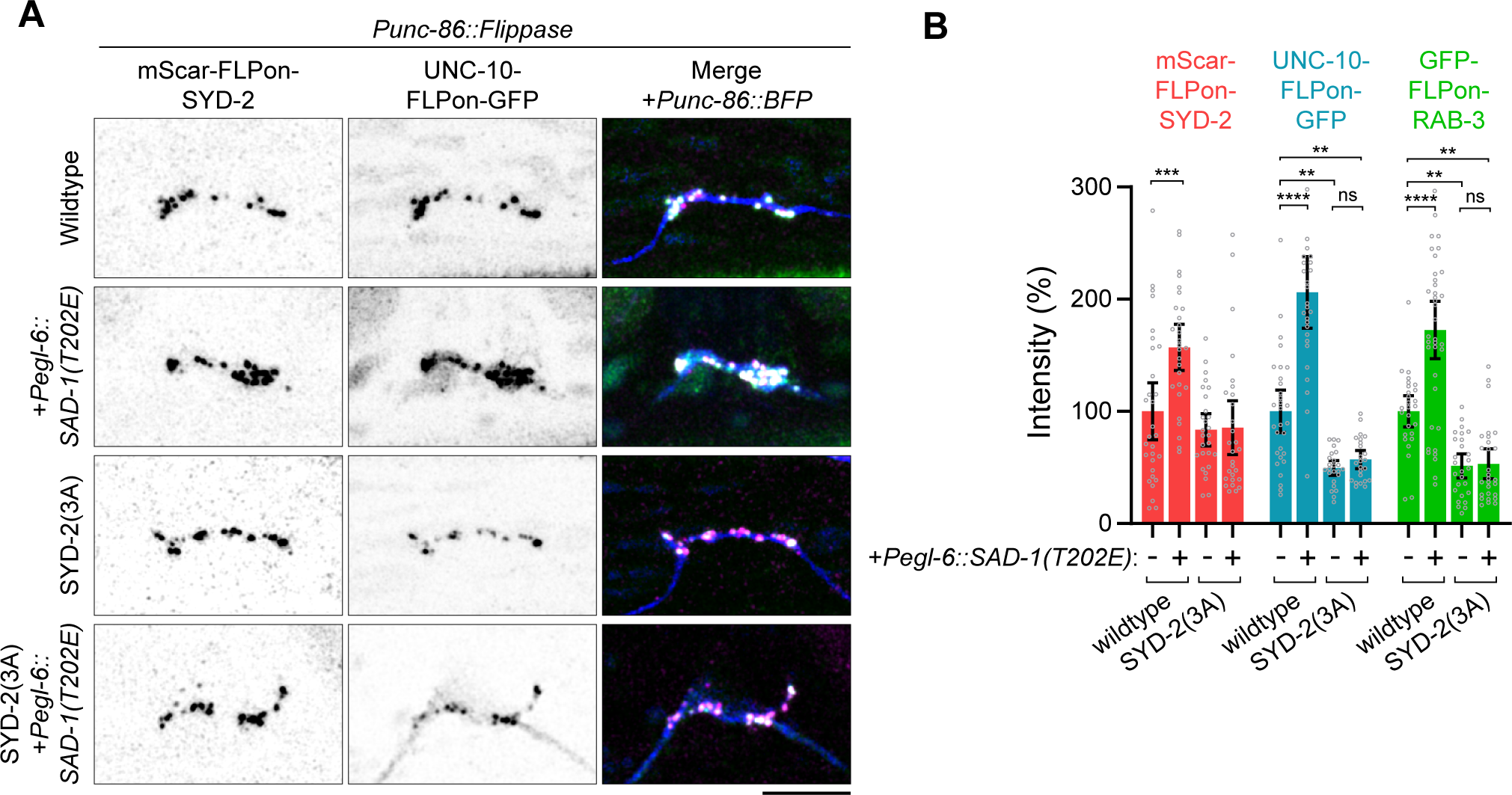
SAD-1 control of active zone assembly requires SYD-2 phosphorylation. A) HSN synapse formation phenotypes visualized with endogenous fluorescent tags in the indicated mutants. *Pegl-6::SAD-1(T202E)* is an overexpression of constitutively active SAD-1 kinase in HSN. Scale bars, 5 µm. B) Quantification of HSN intensities in (A). **, p < .01; ***, p < .001; ****, p < 0.0001; ns, not significant.

#### Phosphorylation of SYD-2 activates phase separation by relieving autoinhibition

In previous examples of phase separation regulation by phosphorylation, direct phosphorylation of intrinsically disordered regions is thought to alter phase separation properties^18^. Here, we find phosphorylation in a folded domain adjacent to an IDR to be responsible for modulating phase separation. We instead hypothesized that these phosphosites may regulate phase separation through conformational changes or intramolecular interactions. To test this *in vitro*, we split the SYD-2 protein in two (Figure 6A), into an N-terminal fragment containing the IDR (Nter), and a C-terminal fragment containing the SAM domains (SAM). We confirmed the Nter fragment was capable of phase separation, as shown previously^9^, and found the SAM fragment alone was not (Figure 6B). The addition of the SAM domains to the Nter condensates results in their incorporation, supporting a possible interaction between the two (Figure 6B). To determine the impact of SAM addition on phase separation, we quantitatively characterized phase diagrams for SYD-2’s Nter alone versus Nter + SAM condensates (Figure 6C). Interestingly, the addition of SAM domains robustly increased the critical concentration of Nter phase separation. Further, FRAP assays to measure internal liquid dynamics of these condensates showed the addition of SAM domains inhibited liquidity (Figure 6D). These data indicate SYD-2’s SAM domains inhibit the protein’s ability to phase separate.

**Figure 6.**
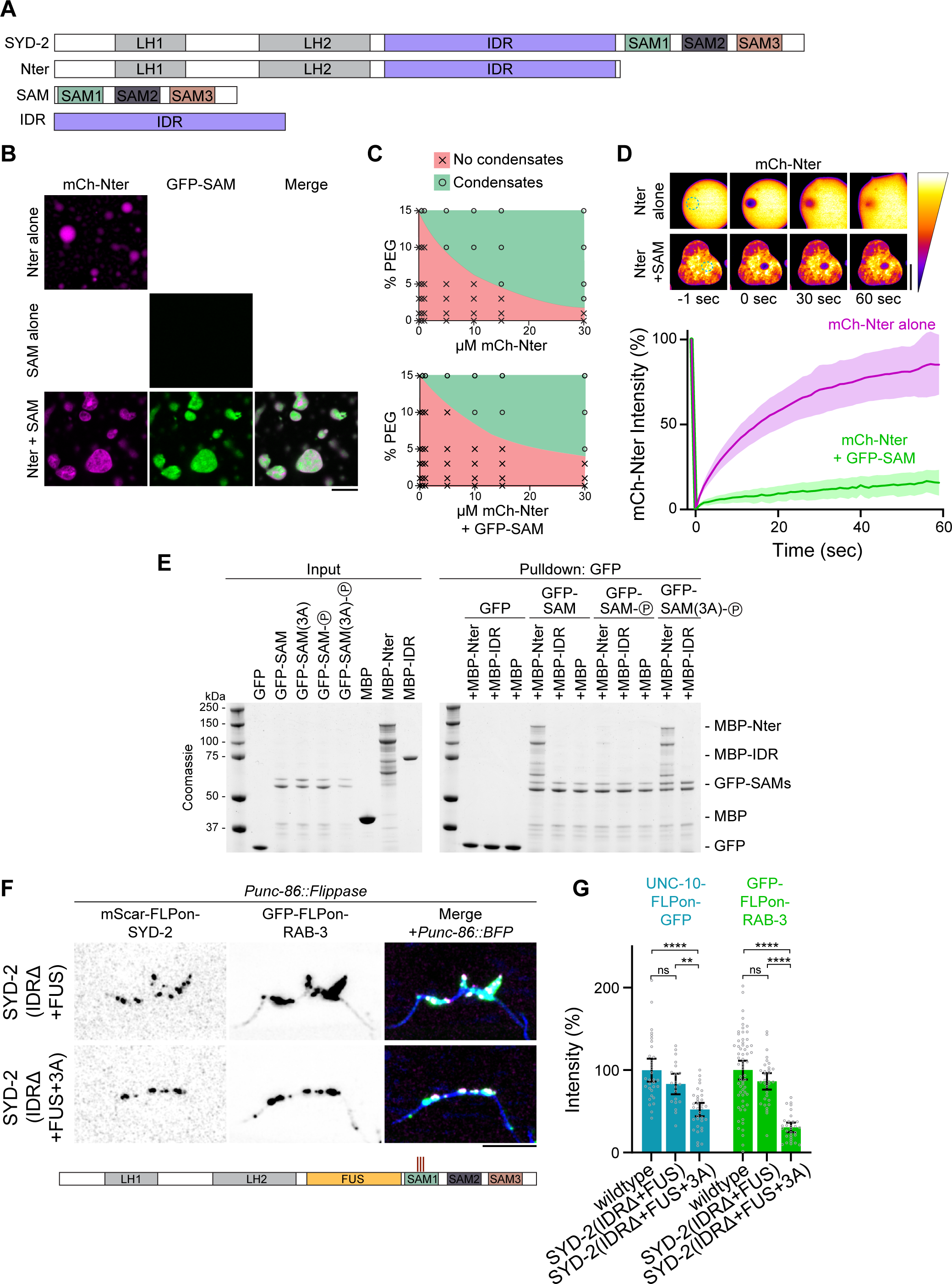
SAD-1 phosphorylation relieves SYD-2 autoinhibition to activate phase separation. A) Domain schematic of SYD-2 and constructs used in this figure. B) *In vitro* phase separation of SYD-2 Nter and SAM domains alone and together. Dilution to 150 mM NaCl and addition of 5% PEG 3000 were used to induce phase separation. C) Quantitative phase diagrams of SYD-2 Nter and Nter+SAM. Addition of SAM domains increased the concentration and crowding conditions required for phase separation, indicating inhibition of Nter phase separation. D) Fluorescent recovery after photobleaching measuring internal dynamics of mCh-SYD-2(Nter) condensates with and without the addition of SAM domains. Quantification below. Addition of SAM domains to SYD-2 Nter condensates inhibits dynamics of in vitro condensates. E) Coomassie gel of *in vitro* binding assay between SYD-2 domains. Constructs labelled “(P)” were pre-phosphorylated by SAD-1 prior to the binding assay. Binding is specific between Nter and SAM domains and is inhibited by phosphorylation of 3 sites (S896, S901, S904). F) SAD-1 phosphorylation continues to regulate phase separation of a FUS replacement construct, where SYD-2’s IDR is replaced by a FUS phase separation motif. Scale bar, 5 µm. G) Quantification of HSN intensity from (F). ****, p < 0.0001; ns, not significant.

These results led us to consider that SYD-2’s SAM domains might bind to its Nter in order to inhibit phase separation. We hypothesized that phosphorylation may release this inhibitory binding to enable phase separation. To test this model, we performed *in vitro* binding assays between SYD-2 N-terminal fragments and SAM domains (Figure 6E). We find that SYD-2’s SAM domains directly interact with an Nter fragment containing the IDR, but an IDR fragment alone is not sufficient for the interaction. We therefore conclude that the SAM domains interact with the coiled-coil regions in SYD-2’s N-terminus. We tested if phosphorylation modulates this interaction by pre-phosphorylating SYD-2’s SAM domains with the SAD-1 kinase. SAD-1 phosphorylation inhibited SAM domain interaction with the N-terminal fragment. This inhibition specifically required the 3 phosphosites we determined functioned in synapse formation (Figure 4), as a SAD-1-phosphorylated SAM(3A) construct restored binding to the N-terminus. Thus, we conclude SYD-2’s SAM domains directly interact with its N-terminus to inhibit phase separation, which is released by specific SAD-1 phosphorylation.

Since phosphorylation relieves SAM-Nter binding and activates phase separation, we tested if the removal of inhibitory SAM domains or if phosphomimetic mutations of SAD-1 phosphosites would precociously activate SYD-2 *in vivo*. However, we find a SAM1-3Δ mutation resulted in a severe loss of SYD-2 protein (Figure S6A,C) and a loss-of-function phenotype (Figure S6D). A phosphomimetic 3E (S896E, S901E, S904E) mutant also drastically decreased SYD-2 levels and resulted in loss-of-function phenotypes (Figure S6B-D). It remains plausible that these modifications precociously activate SYD-2, however the resulting phase separation is clearly not occurring at the proper synaptic localization as when under its native regulation. We can conclude that SYD-2’s SAM domains are critical for protein stability.

Together, our results support a model for intramolecular regulation of IDR-mediated phase separation. SYD-2’s unphosphorylated C-terminal SAM domains bind in the N-terminus, inhibiting the intervening IDR’s phase separation. Intriguingly, this mechanism may be agnostic to the intermediate IDR, as inhibitory binding occurs between two IDR-flanking domains. To test the generalizability of the intramolecular autoinhibition, we introduced the 3A phosphosite mutation into an allele that replaced SYD-2’s IDR with the IDR of FUS (Figure 6D). Swapping SYD-2’s IDR for FUS’s was previously seen to rescue its phase separation and synapse formation functions^9^. Indeed, the addition of the 3A mutation inhibited synapse formation in SYD-2(IDRΔ+FUS), indicating these sites and the intramolecular interaction they control are capable of regulating the phase separation of generic IDR motifs in the protein (Figure 6F-G). This result also agrees with the lack of phenotypes in a 26A IDR mutant (Figure S3). Thus, SAD-1 phosphorylation controls an autoinhibitory interaction between N- and C-terminal domains in SYD-2 that inhibit phase separation of an intermediate IDR.

#### SAD-1 is poised at nascent synapses to activate active zone phase separation

Our data suggest SYD-2 phosphorylation by SAD-1 activates the protein to phase separate and assemble the presynaptic active zone. Indeed, SAD-1 localizes strongly to presynaptic sites, as determined by an endogenous GFP-SAD-1 allele (Figure 2E). To determine if this localization is dependent on SYD-2 and the active zone or an upstream synaptic adhesion molecule, we imaged SAD-1 localization in a *syd-2Δ* mutant. We find that SAD-1 localization does not require *syd-2* and active zone formation: SAD-1 is slightly decreased, but generally remains localized to presynaptic sites when SYD-2 is absent. However, in a *syg-1Δ* mutant, a synaptic cell adhesion molecule responsible for establishing the HSN synaptic region and initiating synapse formation^31, 32^, SAD-1 localization is lost (Figure 7A-B). Therefore, SAD-1 is positioned at presynaptic sites through upstream synaptic cell adhesion cues where it can phosphorylate and activate SYD-2’s phase separation to build the presynaptic active zone (Figure 7C).

**Figure 7.**
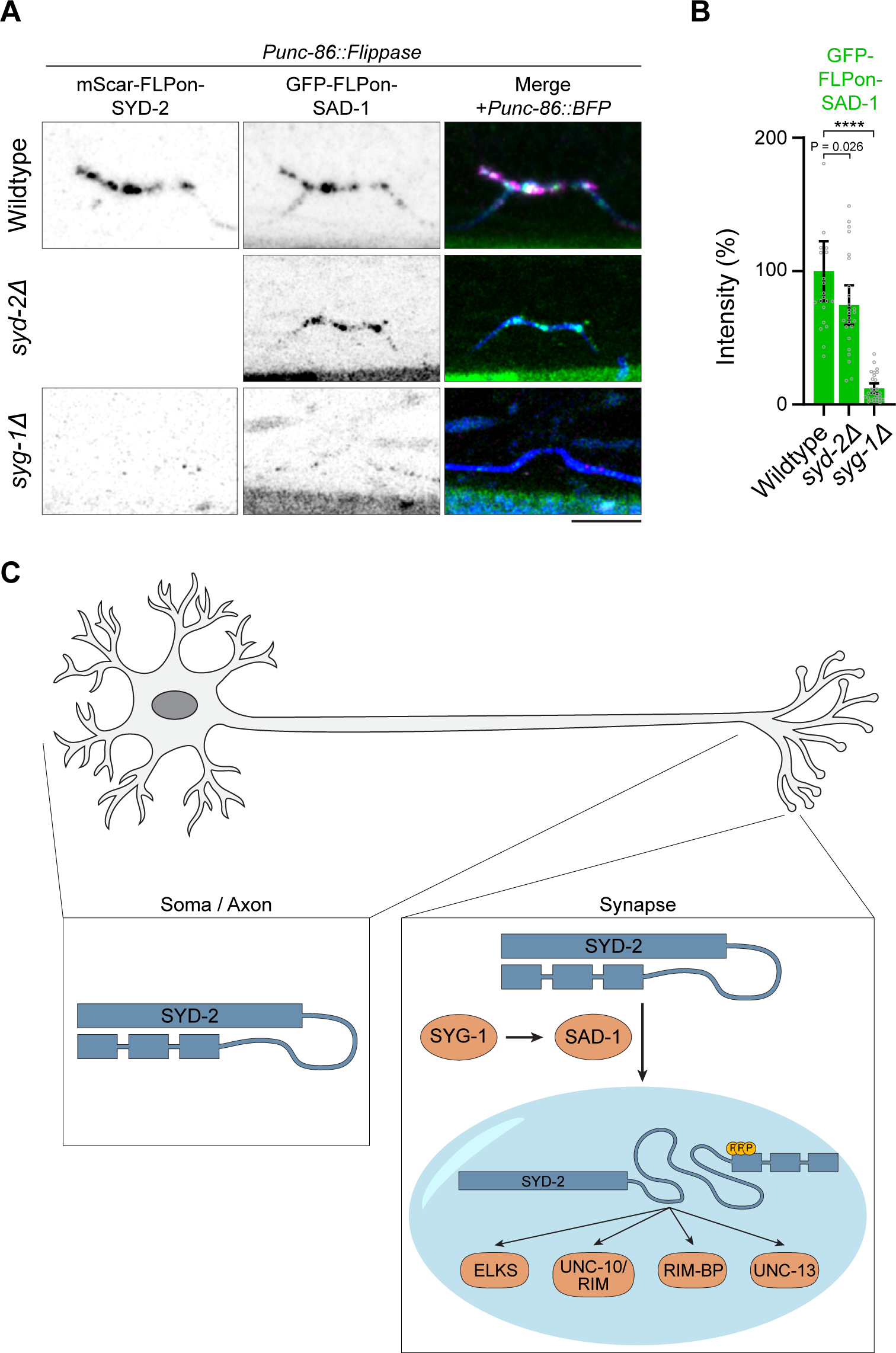
SAD-1 localization to presynapses independent of SYD-2 positions it to activate phase separation and active zone formation. A) Endogenous GFP-SAD-1 localization to HSN synapses with or without *syd-2* or *syg-1*. B) Quantification of GFP-SAD-1 HSN intensity in ****, p < 0.0001. C) Model of SAD-1 activation of SYD-2 phase separation and active zone formation.

## Discussion

In this study, we have identified the SAD-1 kinase to activate presynaptic SYD-2/Liprin-α phase separation and active zone assembly. SAD-1 has been implicated in various neuronal development processes, and here we find specific loss-of-function and overexpression phenotypes for SAD-1 in presynaptic active zone assembly. We identify a key SAD-1 substrate in SYD-2/Liprin-α that accounts for SAD-1’s function in presynaptic assembly. SYD-2 was previously shown to phase separate during presynaptic active zone assembly; we find SAD-1 phosphorylation is a key regulatory mechanism that activates SYD-2’s phase separation function at nascent synapses.

The group of phosphorylation sites we identify as responsible for SYD-2 regulation are well conserved with mammalian Liprin-α homologs of SYD-2. The serine 896, 901, and 904 residues are conserved in all four Liprin-αs, with the exception of an S901A change in Liprin-α3. In addition, phosphorylation has been observed at multiple of these sites in mammalian genome-wide phosphoproteomics datasets^33^. Considering this conservation, the functional conservation of phase separation^10^, and the functional conservation of SYD-2/Liprin-α in active zone assembly in mammals^3^, we consider it likely the regulation discovered here is conserved.

Previously, phosphorylation by PKC was also observed at S760 in Liprin-α3, which was seen to regulate its ability to phase separate. This site was not, however, necessary for Liprin-α2 to phase separate. The S760 site is not explicitly conserved in SYD-2, though it is very near a number of IDR phosphosites we tested and found had no clear function (12A, Figure S3). IDR sequences have been observed to diverge quickly at the sequence level over evolution, while presumably preserving function^34^. It will be necessary to test if other sites in SYD-2 are phosphorylated by PKC to substitute a similar regulation.

A key question in neuronal development is how intracellular assembly of synaptic structures is connected to adhesion molecule-based specificity mechanisms^4^. The SAD-1 activation signal we have identified likely occurs at nascent presynaptic sites where SAD-1 is strongly localized. Previously, SAD-1 has been seen to moderately depend on SYD-2 for localization^22^, placing it “downstream” in an assembly hierarchy. However, these experiments were performed with overexpressed proteins and were not quantitative. With knock-in fluorescent tags, representing endogenous protein levels and localization, we see only a minor dependence of SAD-1 localization on SYD-2; 80% of SAD-1 protein remains localized to presynaptic sites in the absence of SYD-2. We find that SAD-1 localization to presynapses is instead dependent on SYG-1, an upstream synaptic cell adhesion molecule that instructs the location of synapses in HSN and select other neurons^31, 35^, likely through the formation of an actin network^36^. SAD-1 has been seen to bind NAB-1^37^, an actin-binding protein which also depends on SYG-1 for localized to presynaptic sites. Thus, the data suggests upstream synaptic cell adhesion molecule SYG-1 localizes SAD-1 to nascent synapses where it activates SYD-2 by phosphorylation of SAM domains.

In addition to SYD-2, we find a variety of possible SAD-1 substrates in our phosphoproteomics screen (Table S1). Intriguingly, the substrates identified are enriched in proteins involved in signaling (p = 1.5*10^-8^), cytoskeleton (p = 1.8*10^-7^), and neuronal function (p = 1.1*10^-4^). Additional study on these substrates may reveal links to SAD-1’s described functions in neuronal polarity and neurite outgrowth. SYD-2 phosphorylation appears to account for SAD-1’s specific function in presynapse assembly, as a SYD-2(3A) mutant phenocopies a *sad-1Δ* and SAD-1 overexpression is unable to impact synaptic accumulation in SYD-2(3A).

Beyond the neuronal development signaling revealed here, we have identified a novel mechanism that controls phase separation by an internal intrinsically disordered region. In most examples of phase separation regulation by post-translational modifications, direct modification of IDRs cause changes in biochemical properties which impact the ability to phase separate^18^. Here, instead, we find phosphorylation controls an intramolecular binding interaction and conformational change. When dephosphorylated, binding between domains surrounding the IDR prevents its phase separation. Phosphorylation releases this binding, enabling IDR phase separation. This mechanism is particularly intriguing for spatial regulation and activation of phase separation, as the newly translated, unphosphorylated protein will adopt an autoinhibited form. Only upon phosphorylation at the proper place, in this case nascent synaptic sites, is autoinhibition relieved to enable IDR phase separation. As SYD-2 condensates scaffold a variety of additional active zone components^3, 9, 22^, autoinhibition is an attractive mechanism to prevent improper assembly at the wrong place and time in the neuron.

Curiously, this mechanism was agnostic to the IDR within SYD-2 performing the phase separation. How binding between folded domains flanking the IDR actually inhibits its phase separation is not yet clear. It is possible that the bound state “stretches” the IDR into an extended conformation that prevents the multivalent interactions underlying phase separation^5, 6^. As more IDRs are identified to function in condensate formation, similar autoinhibitory and extra-IDR regulation may be at play due to the advantageous autoinhibition, allowing the cell to prevent condensate formation until needed.

## Acknowledgements

This work was funded by the Howard Hughes Medical Institute, a Helen Hay Whitney postdoctoral fellowship to N.A.M., and NIH K99 NS123233 to N.A.M.

## Author Contributions

N.A.M. and K.S. conceptualized the study. N.A.M. carried out the experiments and analyses. L.T. and M-Q.D. performed the *in vivo* phosphoproteomics. N.A.M. wrote the paper. N.A.M., L.T., M-Q.D., and K.S. edited the paper.

## Methods

### *C. elegans* methods

*C. elegans* strains (Table S3) were grown on OP50 *E. coli*-seeded nematode growth media (NGM) plates at 20°C, following standard protocols^38^. N2 Bristol is the wildtype reference strain. ^15^N wildtype worms for phosphoproteomics were grown for at least ten generations on ^15^N-labeled, MG1655 *E. coli*-seeded, nitrogen-free NGM plates at 20°C, as described previously^39^. ^15^N MG1655 was prepared in M9 minimal media containing ^15^NH_4_Cl as the sole nitrogen source.

Nerve ring imaging was performed on comma-stage embryos. HSN imaging was performed on synchronized early L4 hermaphrodites. HSNL (left) was exclusively imaged due to its advantageous separation from the nerve cord. Aldicarb assays were performed on day 1 adult hermaphrodites on NGM plates containing 1 mM Aldicarb (Millipore-Sigma). Arrays were created by gonadal microinjection. *Pegl-6::SAD-1(T202E)* arrays were injected at 30 ng/uL with a 50 ng/uL *Podr-1::GFP* coinjection marker. FLPon tags were constitutively flipped out by temporary expression of a germline *Peft-1::Flippase*.

### Constructs and CRISPR-Cas9 genome editing

Constructs (Table S4) were created with an isothermal assembly (Gibson) method^40^. pNM171 *Punc-86::tagBFP2-SL2-FLP* was assembled into an empty pSK vector from the 5,102-nucleotide promoter of *unc-86*, a *C. elegans* codon-optimized tagBFP2 (pJJR81, Addgene #75029), an SL2 site, Flippase (pDML63^41^), a self-excising Hygromycin resistance cassette^42^, and flanking 500-bp homology arms to the MosSCI site ttTi5605^43^. pNM172 *Pegl-6::SAD-1(T202E)* was assembled into a pSM delta vector from the 3,527-nucleotide promoter of *egl-6* and SAD-1 cDNA. A T202E activating mutation was introduced with site-directed mutagenesis, designed based on alignment to MAP kinase activation loops^44, 45^. Lambda phosphatase from *Escherichia* phage lambda was synthesized as a *C. elegans* codon-optimized gBlock (IDT) and assembled into a pSK vector. A H76N inactivating mutation^46^ was introduced with site-directed mutagenesis. SYD-2 SAM(844-1139), SYD-2 IDR(517-843), and SAD-1 constructs were assembled into pHis6-GFP, pHis6-mCherry, and pMBP-his empty vectors. 3A, 3E, Bottom4A, and Tail3A phosphomutants were introduced into SYD-2 constructs using site-directed mutagenesis. 10A, 12A, and 26A phosphomutants were synthesized as gBlocks (IDT) and assembled into pMBP-his vectors.

Endogenous genome modifications were created as previously described^9, 47^, through gonadal microinjection of Cas9 (IDT), tracrRNA (IDT), gRNA (IDT), and PCR repair template mixtures. mScarlet-I-FLPon cassettes used FRT(F3) sites (GAAGTTCCTATTCTTCAAATAGTATAGGAACTTC) and GFP-FLPon cassettes used FRT (GAAGTTCCTATTCTCTAGAAAGTATAGGAACTTC) sites for simultaneous compatibility. WySi974, a single-copy *Punc-86::tagBFP2-SL2-FLP* that drives Flippase and a tagBFP2 morphology marker in HSN, was created by insertion into the MosSCI ttTi5605 site on chromosome V as a convenient and well-established expression locus^43^ using a hygromycin resistance strategy^42^. SYD-2 phosphomutants were introduced into SYD-2(IDRΔ) or SYD-2(SAM1-3Δ) strains with repair templates generated by PCR from pSK vectors or gBlocks, described above. The IDRΔ+FUS replacement was created in an mScarlet-I-FLPon-SYD-2 strain as previously described^9^. The following gRNAs were used for genome edits: mScar-FLPon-SYD-2: AGAAATATGAGCTACAGCAA; UNC-10-FLPon-GFP: GATTCCGATGTATCAGTTGG; Lambda phosphatase (wildtype and H76N): TTTAATTTAACTAACTAACT; IDRΔ: GAACTGCGCAATTCCAGTCA and GGCGAGCAGTCGGGCACAGA; GFP-FLPon-SAD-1: CATGACTGCGCTCGTCAATC; SAMΔ: TCCAACTGTTGTTGCCTGGC and TTTAATTTAACTAACTAACT; SAM phosphomutants (3A, Bottom4A, Tail3A, 10A, 3E): AATTACCAAGCAACAACAGT; IDR phosphomutants (12A, 26A): AATGCAAGAACTGCGCAATT. All genome-edited strains were outcrossed and verified by sequencing.

### Protein methods

*C. elegans* were prepared for immunoprecipitation by washing 3x in M9 buffer, resuspension in 1 volume of TBS containing 1 mM PMSF and cOmplete protease inhibitor cocktail (Roche), and dropwise addition to liquid nitrogen to snap freeze. Frozen droplets were ground to a fine powder with a mortar and pestle and resuspended in a RIPA buffer with phosphatase inhibitors (50 mM Tris pH 7.4, 150 mM NaCl, 1% NP-40, 0.1% SDS, 0.1% sodium deoxycholate, 10 mM EDTA, 1 mM EGTA, 1 mM Na_3_VO_4_, 60 mM β-glycerophosphate, 2.5 mM sodium pyrophosphate, 50 mM sodium fluoride, 2 mM benzamidine, 1 mM PMSF, and cOmplete protease inhibitor cocktail). Lysates were sonicated to complete worm lysis and cleared by centrifugation at 60k xg. GFP was pulled down with GFP-trap agarose beads (ChromoTek) and mScarlet was pulled down with RFP-trap agarose beads (ChromoTek). Select samples were treated on-bead with lambda protein phosphatase (NEB) in the manufacturer’s PMP buffer to remove phosphorylation. Immunoprecipitated sample was eluted in SDS sample buffer, run on a 4-12% Bis-tris gel (Invitrogen), transferred to a PVDF membrane, and blotted for GFP (Roche 11814460001), mScarlet (ChromoTek 6G6), or phospho-serine/threonine (Abcam ab17464).

*C. elegans* were prepared for phosphoproteomics as described previously^39^. ^14^N *sad-1Δ* and ^15^N-labeled wildtype worms were mixed at a 1:1 volume ratio. The sample mixtures were lysed by resuspension in 2x RIPA with 2x EDTA-free proteinase inhibitor cocktail (Roche) and 2x PhosSTOP (Roche). Samples were homogenized with a FastPrep-24 (MP Biomedicals) and cleared by centrifugation at 20,000 xg for 30 min. Proteins were precipitated from cleared samples with acetone and resuspended in a urea solution (8 M urea, 100 mM Tris-HCL pH 8.5) for trypsin digestion.

Purified protein constructs were produced in Rosetta2(DE3) *E. coli* cells grown at 37 °C in TB medium and induced with 0.4 mM IPTG overnight at 18 °C. Cells were lysed in 20 mM Tris pH 7.4, 500 mM NaCl (high salt to inhibit phase separation), 5 mM β-mercaptoethanol, 1 mM phenylmethylsulfonyl fluoride (PMSF), and cOmplete protease inhibitors (Roche) with 0.2 mg/mL lysozyme. Lysates were spun at 30,000 xg to clear and remove pre-condensed or aggregated protein. His6-GFP and His6-mCherry proteins were purified with cOmplete His tag resin (Roche) according to the manufacturers protocols as previously described^9^. pMBP-his proteins containing MBP on their N-terminus and 8xHis on their C-terminus were purified in a two-step process to select for full-length products. First, lysates were incubated with amylose resin (New England Biolabs) for 2 hours, drained in a polyprep column (Biorad), and washed with 12 volumes of wash buffer (20 mM Tris pH 7.4, 500 mM NaCl, 5 mM β-mercaptoethanol). Protein was eluted with wash buffer containing 10 mM maltose and subsequently bound to c0mplete His tag resin (Roche) for 2 hours. The resin was washed 3x to remove free MBP and truncated proteins. Full-length protein was eluted with wash buffer + 250 mM imidazole. Samples were dialyzed overnight with 10,000 MWCO SnakeSkin dialysis tubing (Thermo), concentrated to roughly 50-100 µM with 10,000 MWCO Amicon Ultra centrifugal filters, and snap frozen in aliquots at −80 °C.

*In vitro* kinase assays (modified from ^28^) consisted of 1 µg substrate, 1 µg SAD-1 kinase, 0.1 µg LKB-1 complex (Sigma-Aldrich), 500 µM ATP, and 2 µCi γ^32^P-ATP (Perkin-Elmer) in 25 mM Tris pH 7.4, 10 mM magnesium acetate, 1 mM DTT. SAD-1 and LKB-1 were preincubated to activate the kinase in the absence of γ^32^P-ATP for 30 minutes at room temperature, followed by addition of the substrate and γ^32^P-ATP for an additional 30 minutes at room temperature. Reactions were run on a 4-12% Bis-tris gel and stained with Coomassie SimplyBlue (Invitrogen) before drying. Dried gels were assembled in cassettes with a BAS-IP Phosphor screen (Cytiva) overnight and imaged on an Amersham Typhoon system (Cytiva) with 635 nm excitation.

*In vitro* liquid-liquid phase separation assays were performed as previously described^9^. Purified recombinant proteins were diluted to physiological salt conditions (20 mM Tris pH 7.4, 150 mM NaCl) in the presence of 5% PEG 3000. Phase diagrams were determined on the basis of proteins’ ability to form droplets within 5 minutes in each condition.

The *in vitro* binding assay was performed by binding 1 µg of each purified recombinant GFP construct to GFP-trap agarose beads in 20 mM Tris pH 7.4, 500 mM NaCl, 5 mM β-mercaptoethanol. Select GFP samples were pre-phosphorylated by the SAD-1 kinase as described above (excluding γ^32^P-ATP). 1 µg of each MBP-conjugated prey protein was incubated with the GFP-SAM-bound beads for 1 hour. Beads were washed and eluted with SDS sample buffer, and bound protein was visualized with Coomassie-stained SDS-PAGE.

### Phosphoproteomics

Samples were prepared for mass spectrometry as previously described^39^. 10 mg of protein (^14^N and ^15^N mixtures) was reduced with 5 mM TCEP, alkylated with 10 mM iodoacetamide, and digested with trypsin overnight at 37°C. Digested peptides were separated into 12 fractions on a Xtimate C18 reverse phase HPLC column (10 x 250 mm, 5 µm, Welch Materials) with an Agilent 1200 Series HPLC. Each fraction was acidified and enriched for phosphopeptides using a PolyMAC-Ti Enrichment Kit (Tymora Analytical). The resulting phosphopeptides were resolved in 0.25% formic acid buffer for mass spectrometry analysis.

Each fraction was analyzed in two technical replicates by a Q-Exactive mass spectrometer (Thermo Fisher Scientific) interfaced with an Easy-nLC1000 reverse phase chromatography system (Thermo Fisher Scientific). Data acquisition, phosphopeptide identification, and quantification was performed exactly as previously described^39^. A wildtype dataset from this study (^14^N wildtype mixed with ^15^N wildtype sample) was used to normalize for altered phosphoproteomes in ^15^N samples. GO enrichment analysis was performed with WormCat^48^.

*In vitro* kinase assay phosphosite mapping was performed by MS Bioworks (Ann Arbor, MI). Phosphorylated SYD-2 constructs in gel slices were reduced with 10 mM dithiothreitol (DTT), alkylated with 50 mM iodoacetamide, and digested with either trypsin, chymotrypsin, or elastase (Promega). Samples were analyzed by nano LC-MS/MS on a Waters M-Class HPLC coupled to a ThermoFisher Orbitrap Fusion Lumos mass spectrometer. Peptides were loaded on a trapping column and eluted over a 75 µm analytical column at 350 nL/min. Both columns were packed with Luna C18 resin (Phenomenex). The mass spectrometer was operated in data-dependent mode, with the Orbitrap operating at 60,000 FWHM and 15,000 FWHM for MS and MS/MS respectively. APD was enabled and the instrument was run with a 3 sec cycle for MS and MS/MS.

### Microscopy

Fluorescence imaging was performed on a Zeiss LSM 980 Airyscan 2 system with a 63X Plan-Apochromat 1.4NA objective and 405 nm, 488 nm, or 561 nm lasers. The airyscan 4Y multiplex mode was used to capture super-resolution images of HSN synapses, while the 8CO confocal mode on the Airyscan detector was used for embryonic synapse FRAP to maximize sensitivity. *In vitro* condensate FRAP was performed with the confocal LSM detector. Animals were imaged live on 5% agarose pads in 1 mM levamisole.

### Image analysis

Airyscan images were processed with Zeiss ZEN software. Image analysis was performed in ImageJ/Fiji (National Institutes of Health). FRAP curves were calculated after background and time-course photobleaching correction and normalized to 100% for visualization. HSN intensity quantification was performed on background-subtracted sum projections of Z-stacks. Representative images are maximum intensity projections that have been cropped and rotated where necessary. Images have inverted greyscale or “fire” Fiji lookup tables applied for visualization.

### Statistics and reproducibility

Statistical comparisons were performed with one-way ANOVA tests and multiplicity-corrected P-values were calculated versus wildtype using Dunnett’s test in Prism 9 (GraphPad). Bar graphs depict all data points, means, and 95% confidence interval error bars. FRAP graphs depict 95% confidence intervals and one-phase association least-squares fits from GraphPad Prism 9. Each *C. elegans in vivo* measurement (FRAP or fluorescence intensity) is a biological replicate taken from a distinct animal. All *in vivo* imaging data were replicated in at least two independent sessions. All *in vitro* experiments were replicated at least twice. Three biological replicates were performed for SAD-1 phosphoproteomics.

---

**Figure S2.**
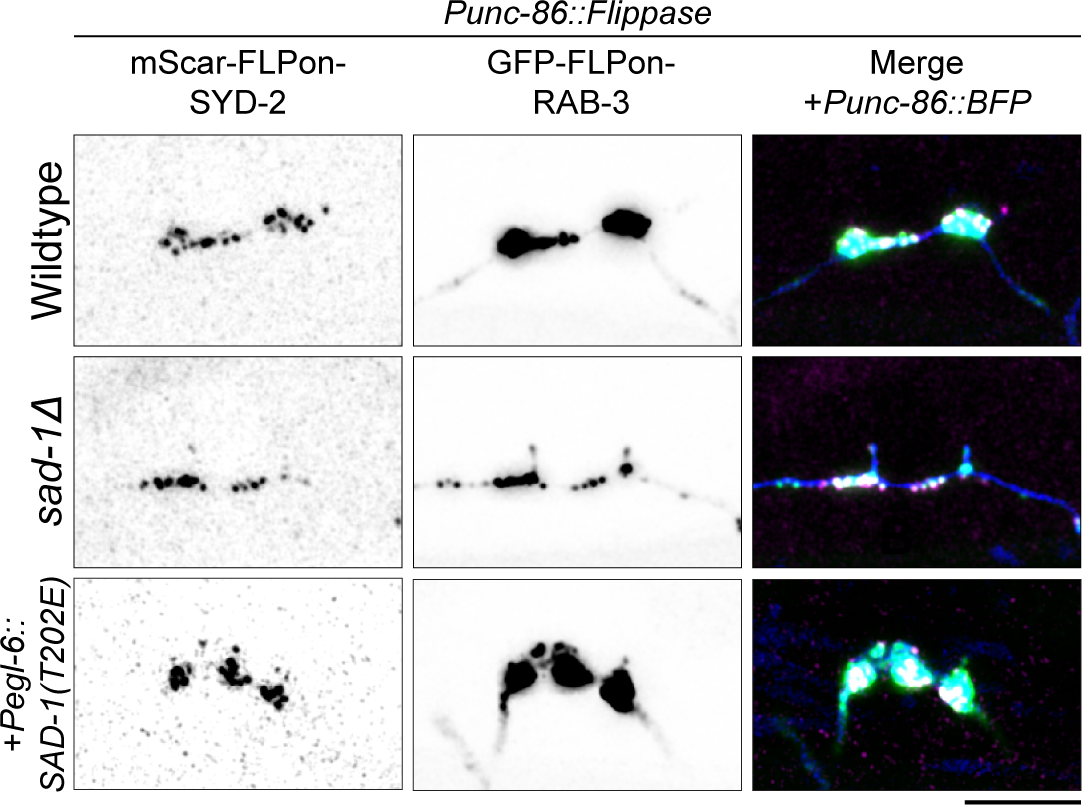
SAD-1 kinase regulation of synaptic vesicle clustering. Synapse formation phenotypes visualized with endogenous GFP-RAB-3 in the indicated mutants. Scale bars, 5 µm.

**Figure S3.**
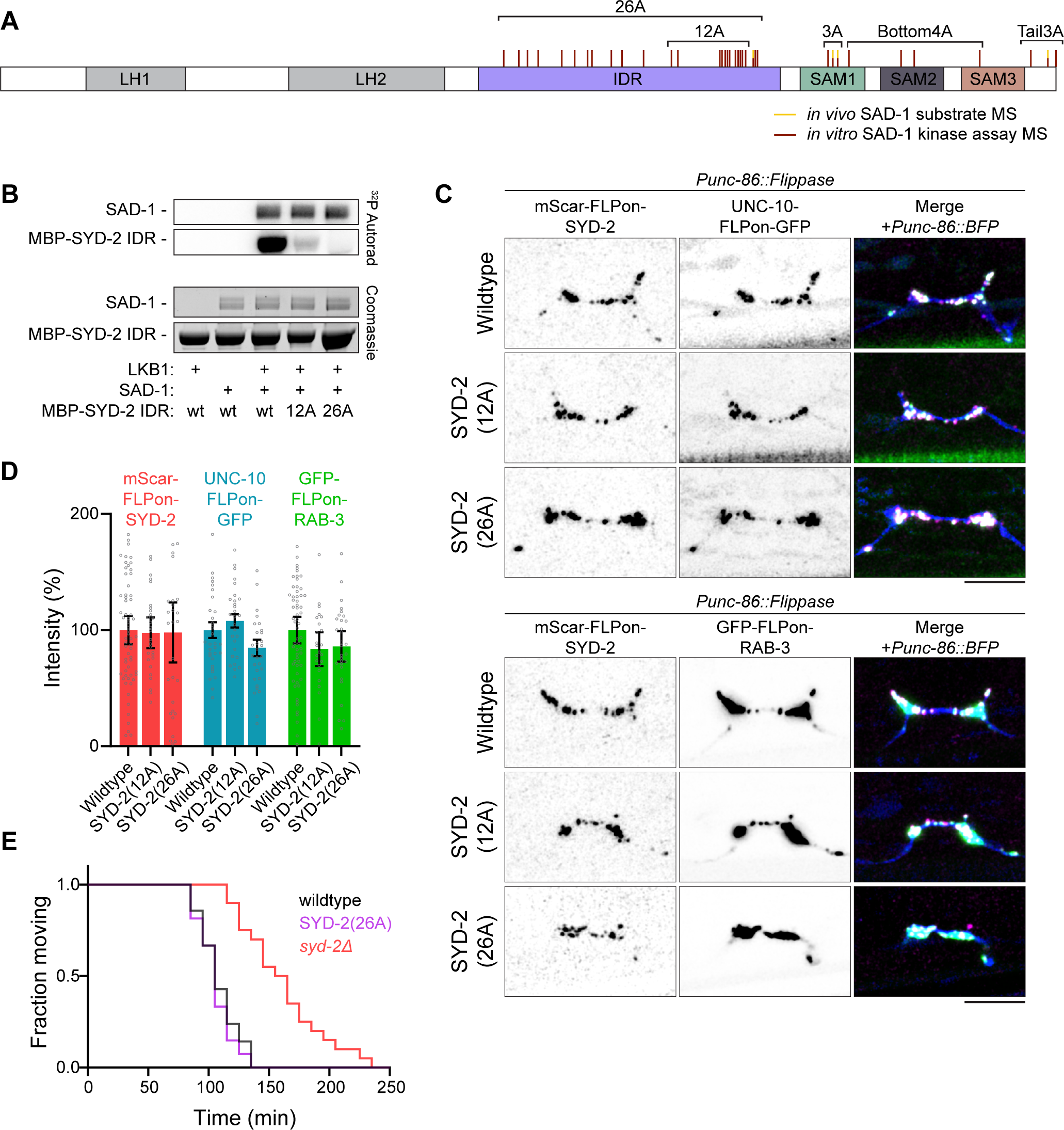
SAD-1 phosphorylation of SYD-2’s IDR *in vitro* is dispensable for *in vivo* function. A) SYD-2 phosphosites identified from *in vivo* phosphoproteomics (yellow) or *in vitro* kinase assays (burgundy). Sites are grouped based on location for follow-up testing. See Tables S1-2. B) *In vitro* kinase assay between SAD-1 and SYD-2 IDR with or without phosphosite mutations. SAD-1 is activated by the LKB-1 kinase complex. C) HSN synapse formation phenotypes visualized with endogenous fluorescent tags in the indicated mutants. Scale bars, 5 µm. D) Quantification of HSN intensities in (C). No significant difference in synapse formation was seen in IDR phosphomutants. Wildtype data is from Figure 1F. E) Aldicarb synaptic transmission assay shows no defects in a SYD-2 IDR phosphomutant. Additional time to paralysis on 1 mM Aldicarb indicates defective synaptic transmission. n > 20 for each genotype.

**Figure S4.**
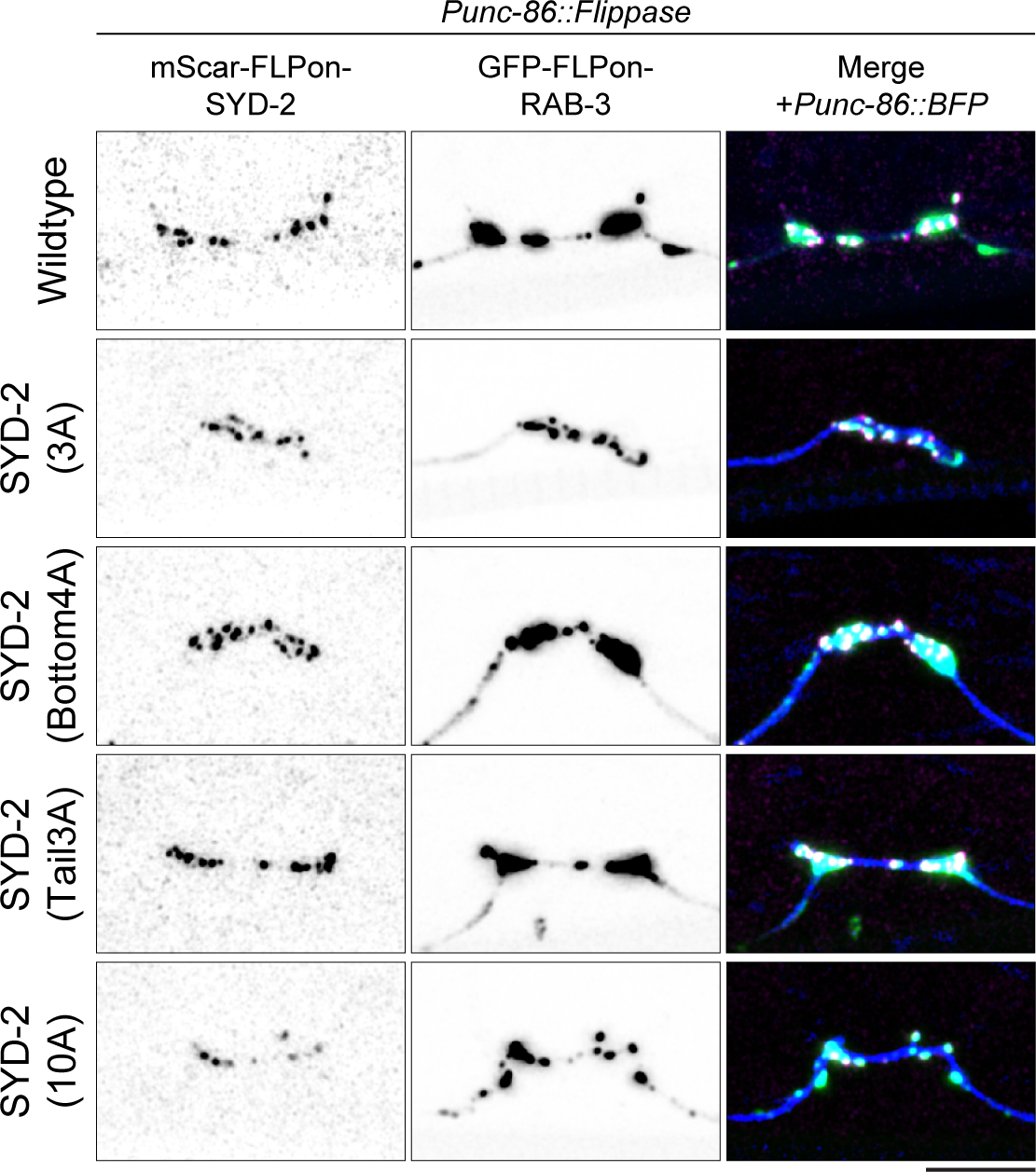
SYD-2 phosphorylation and synaptic vesicles clustering. HSN synapse formation phenotypes visualized with endogenous fluorescent tags in the indicated mutants. Scale bars, 5 µm. Quantification presented in Figure 4B.

**Figure S6.**
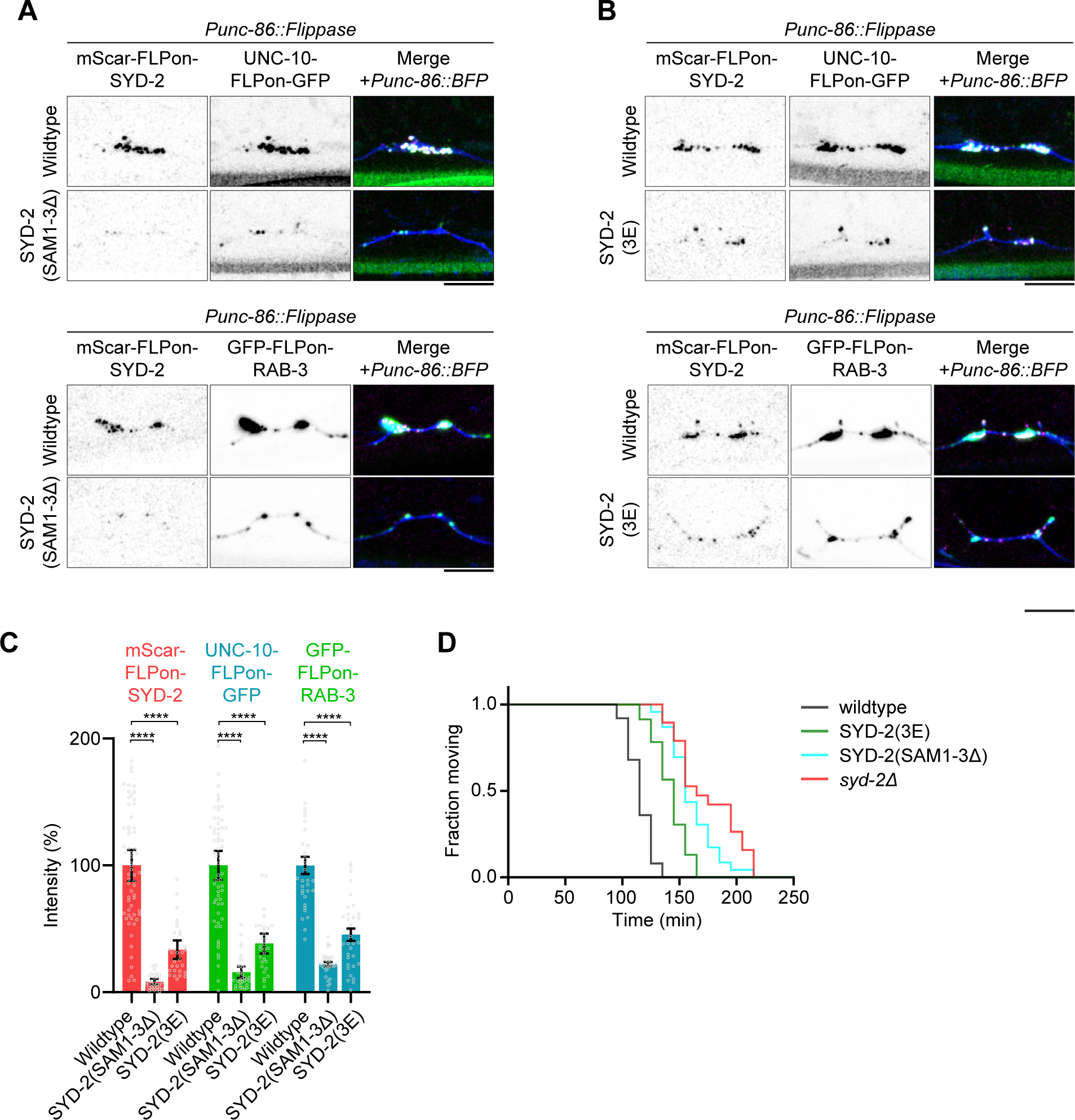
Precocious activation of SYD-2 phase separation renders the protein unstable and unable to reach synaptic sites. A-B) HSN synapse formation phenotypes visualized with endogenous fluorescent tags in the indicated mutants. Removal of SAM domains in (A) or mimicking SAD-1 phosphorylation in (B) leads to reduced SYD-2 at HSN synapses and a subsequent active zone assembly phenotype. C) Quantification of HSN intensities in (A-B). ****, p < 0.0001. D) Aldicarb synaptic transmission assay. Additional time to paralysis on 1 mM Aldicarb indicates defective synaptic transmission. n > 20 for each genotype.

## Notes

### Competing Interest Statement

The authors have declared no competing interest.

## References

1. Südhof, T. C. The cell biology of synapse formation. J. Cell Biol. 220, (2021).

2. Südhof, T. C. The Presynaptic Active Zone. Neuron 75, 11–25 (2012).

3. Emperador-Melero, J. & Kaeser, P. S. Assembly of the presynaptic active zone. Curr. Opin. Neurobiol. 63, 95–103 (2020).

4. Südhof, T. C. Molecular Neuroscience in the 21st Century: A Personal Perspective. Neuron 96, 536–541 (2017).

5. Hyman, A. A., Weber, C. A. & Jülicher, F. Liquid-Liquid Phase Separation in Biology. Annu. Rev. Cell Dev. Biol. 30, 39–58 (2014).

6. Boeynaems, S. et al. Protein Phase Separation: A New Phase in Cell Biology. Trends Cell Biol. 28, 420–435 (2018).

7. Wu, X. et al. RIM and RIM-BP Form Presynaptic Active-Zone-like Condensates via Phase Separation. Mol. Cell 73, 971–984.e5 (2019).

8. Wu, X. et al. Vesicle Tethering on the Surface of Phase-Separated Active Zone Condensates. Mol. Cell 81, 13–24.e7 (2021).

9. McDonald, N. A., Fetter, R. D. & Shen, K. Assembly of synaptic active zones requires phase separation of scaffold molecules. Nature 588, 454–458 (2020).

10. Emperador-Melero, J. et al. PKC-phosphorylation of Liprin-α3 triggers phase separation and controls presynaptic active zone structure. Nat. Commun. 12, 3057 (2021).

11. Milovanovic, D., Wu, Y., Bian, X. & De Camilli, P. A liquid phase of synapsin and lipid vesicles. Science 361, 604–607 (2018).

12. Park, D. et al. Cooperative function of synaptophysin and synapsin in the generation of synaptic vesicle-like clusters in non-neuronal cells. Nat. Commun. 12, 263 (2021).

13. Imoto, Y. et al. Dynamin is primed at endocytic sites for ultrafast endocytosis. Neuron 110, 2815–2835 (2022).

14. Zeng, M. et al. Phase Transition in Postsynaptic Densities Underlies Formation of Synaptic Complexes and Synaptic Plasticity. Cell 166, 1163–1175.e12 (2016).

15. Xing, G. et al. Membraneless condensates by Rapsn phase separation as a platform for neuromuscular junction formation. Neuron 109, 1963–1978.e5 (2021).

16. Wu, X., Cai, Q., Feng, Z. & Zhang, M. Liquid-Liquid Phase Separation in Neuronal Development and Synaptic Signaling. Dev. Cell 55, 18–29 (2020).

17. Wu, X., Qiu, H. & Zhang, M. Interactions between membraneless condensates and membranous organelles at the presynapse: a phase separation view of synaptic vesicle cycle. J. Mol. Biol. 435, 167629 (2022).

18. Li, J. et al. Post-translational modifications in liquid-liquid phase separation: a comprehensive review. Mol. Biomed. 3, 13 (2022).

19. Zielinska, D. F., Gnad, F., Jedrusik-Bode, M., Wiśniewski, J. R. & Mann, M. Caenorhabditis elegans Has a Phosphoproteome Atypical for Metazoans That Is Enriched in Developmental and Sex Determination Proteins. J. Proteome Res. 8, 4039–4049 (2009).

20. Collins, K. M. et al. Activity of the C. elegans egg-laying behavior circuit is controlled by competing activation and feedback inhibition. eLife 5, e21126 (2016).

21. Schafer, W. R. Genetics of Egg-Laying in Worms. Annu. Rev. Genet. 40, 487–509 (2006).

22. Patel, M. R. et al. Hierarchical assembly of presynaptic components in defined C. elegans synapses. Nat. Neurosci. 9, 1488–1498 (2006).

23. Schwartz, M. L. & Jorgensen, E. M. SapTrap, a Toolkit for High-Throughput CRISPR/Cas9 Gene Modification in *Caenorhabditis elegans*. Genetics 202, 1277–1288 (2016).

24. Jiang, X., Sando, R. & Südhof, T. C. Multiple signaling pathways are essential for synapse formation induced by synaptic adhesion molecules. Proc. Natl. Acad. Sci. 118, e2000173118 (2021).

25. Kishi, M., Pan, Y. A., Crump, J. G. & Sanes, J. R. Mammalian SAD Kinases Are Required for Neuronal Polarization. Science 307, 929–932 (2005).

26. Lilley, B. N. et al. SAD kinases control the maturation of nerve terminals in the mammalian peripheral and central nervous systems. Proc. Natl. Acad. Sci. 111, 1138–1143 (2014).

27. Inoue, E. et al. SAD: A Presynaptic Kinase Associated with Synaptic Vesicles and the Active Zone Cytomatrix that Regulates Neurotransmitter Release. Neuron 50, 261–275 (2006).

28. Kim, J. S. et al. A chemical-genetic strategy reveals distinct temporal requirements for SAD-1 kinase in neuronal polarization and synapse formation. Neural Develop. 3, 14 (2008).

29. Crump, J. G., Zhen, M., Jin, Y. & Bargmann, C. I. The SAD-1 Kinase Regulates Presynaptic Vesicle Clustering and Axon Termination. Neuron 29, 115–129 (2001).

30. Kim, J. S. M., Hung, W., Narbonne, P., Roy, R. & Zhen, M. C. elegans STRAD and SAD cooperatively regulate neuronal polarity and synaptic organization. Development 137, 93– 102 (2010).

31. Shen, K. & Bargmann, C. I. The Immunoglobulin Superfamily Protein SYG-1 Determines the Location of Specific Synapses in C. elegans. Cell 112, 619–630 (2003).

32. Shen, K., Fetter, R. D. & Bargmann, C. I. Synaptic Specificity Is Generated by the Synaptic Guidepost Protein SYG-2 and Its Receptor, SYG-. Cell 116, 869–881 (2004).

33. Tsai, C.-F. et al. Large-scale determination of absolute phosphorylation stoichiometries in human cells by motif-targeting quantitative proteomics. Nat. Commun. 6, 6622 (2015).

34. Brown, C. J., Johnson, A. K., Dunker, A. K. & Daughdrill, G. W. Evolution and disorder. Curr. Opin. Struct. Biol. 21, 441–446 (2011).

35. Sengupta, T. et al. Differential adhesion regulates neurite placement via a retrograde zippering mechanism. eLife 10, e71171 (2021).

36. Chia, P. H., Chen, B., Li, P., Rosen, M. K. & Shen, K. Local F-actin Network Links Synapse Formation and Axon Branching. Cell 156, 208–220 (2014).

37. Hung, W., Hwang, C., Po, M. D. & Zhen, M. Neuronal polarity is regulated by a direct interaction between a scaffolding protein, Neurabin, and a presynaptic SAD-1 kinase in Caenorhabditis elegans. Development 134, 237–249 (2007).

38. Brenner, S. The Genetics of Caenorhabditis Elegans. Genetics 77, 71–94 (1974).

39. Li, W.-J. et al. Insulin signaling regulates longevity through protein phosphorylation in Caenorhabditis elegans. Nat. Commun. 12, 4568 (2021).

40. Gibson, D. G. et al. Enzymatic assembly of DNA molecules up to several hundred kilobases. Nat. Methods 6, 343–345 (2009).

41. Lipton, D. M., Maeder, C. I. & Shen, K. Rapid Assembly of Presynaptic Materials behind the Growth Cone in Dopaminergic Neurons Is Mediated by Precise Regulation of Axonal Transport. Cell Rep. 24, 2709–2722 (2018).

42. Dickinson, D. J., Pani, A. M., Heppert, J. K., Higgins, C. D. & Goldstein, B. Streamlined Genome Engineering with a Self-Excising Drug Selection Cassette. Genetics 200, 1035–1049 (2015).

43. Frøkjær-Jensen, C., Davis, M. W., Ailion, M. & Jorgensen, E. M. Improved Mos1-mediated transgenesis in C. elegans. Nat. Methods 9, 117–118 (2012).

44. Nolen, B., Taylor, S. & Ghosh, G. Regulation of Protein Kinases: Controlling Activity through Activation Segment Conformation. Mol. Cell 15, 661–675 (2004).

45. Wu, J.-X. et al. Structural insight into the mechanism of synergistic autoinhibition of SAD kinases. Nat. Commun. 6, 8953 (2015).

46. Zhuo, S., Clemens, J. C., Stone, R. L. & Dixon, J. E. Mutational analysis of a Ser/Thr phosphatase. Identification of residues important in phosphoesterase substrate binding and catalysis. J. Biol. Chem. 269, 26234–26238 (1994).

47. Ghanta, K. S. & Mello, C. C. Melting dsDNA Donor Molecules Greatly Improves Precision Genome Editing in Caenorhabditis elegans. Genetics 216, 643–650 (2020).

48. Holdorf, A. D. et al. WormCat: An Online Tool for Annotation and Visualization of Caenorhabditis elegans Genome-Scale Data. Genetics 214, 279–294 (2020).

